# Real-time tagging of biomedical entities

**DOI:** 10.1101/078469

**Authors:** Evangelos Pafilis, Lars Juhl Jensen

## Abstract

Automatic annotation of text is an important complement to manual annotation, because the latter is highly labor intensive. We have developed a fast dictionary-based named entity recognition system, which is used for both real-time and bulk processing of text in a variety of biomedical web resources. We propose to adapt the system to make it interoperable with the PubAnnotation and Open Annotation standards.

## Software implementation

The core of our named entity recognition server is a highly optimized dictionary-based tagging engine implemented in C++. The core tagger is able to process in the order of thousands of PubMed abstracts per second with a single CPU thread and is inherently thread-safe, allowing for perfect scalability in multi-threaded use. It is also available as a Python module, which has been generated in part by the Simplified Wrapper and Interface Generator (SWIG).

To make the tagger available as as a web service, we have developed a multi-threaded Python HTTP server that utilizes the tagger Python module. To ensure efficiency and robustness, tagging requests are processed by a thread pool via a priority queue. If a user already has many requests in the queue, further requests are rejected with an error code to prevent a single user from blocking the service. Requests are also rejected if the document size exceeds 10 MB.

## Applications of the tagger

The tagger has already been applied in a variety of different ways. Its high speed makes it well suited for real-time text mining of web pages, which we have utilized in the augmented browsing tool Reflect (1, 2) and the interactive annotation tool EXTRACT (3).

The same tagging engine forms the basis for several command-line tools. We have previously published open-source taggers for named entity recognition of organisms (4) and environments (5) in large text corpora. Such taggers, combined with other dictionaries and a comention-based scoring scheme, also form the basis for extraction of associations among genes/proteins (6), small molecule compounds (7), cellular components (8), tissues (9), and diseases (10). These text-mining results are normalized to identifiers from suitable databases (11–14) and ontologies (15–18), integrated with associations from other data sources, and made available as a suite of web resources (4–10). The tagging results are also available as JSON-based Linked Data (JSON-LD) via a RESTful Open Annotation API described below (19).

## RESTful web services

The real-time tagger can be accessed via standard HTTP requests with the following syntax: http://tagger.jensenlab.org/{method}?document={text}&entity_types={types}&…, where {`method`} is either GetEntities or GetHTML, {`text`} is the plain or HTML-formatted text to be processed, and {`types`} specifies the types of entities to be tagged. These and additional optional parameters are explained in more detail in Table 1.

**Table 1.**
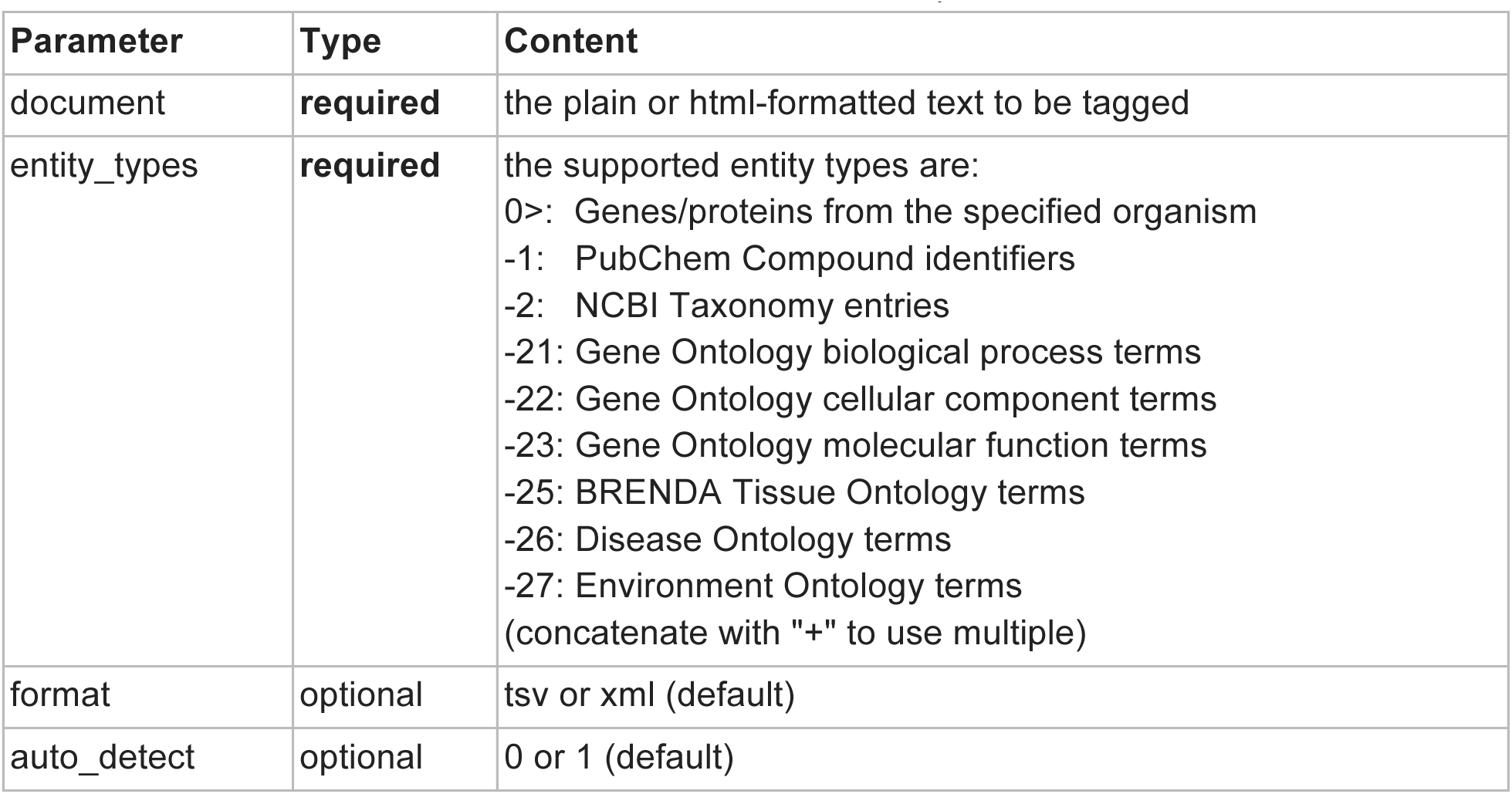
Parameters for the `GetEntities` and GetHTML requests.

The `GetEntities` and `GetHTML` methods both tag the specified types of entities within the provided text document. However, they differ in the results that are returned. `GetEntities` returns the unique list of the entities identified in the document, either in tab-delimited or in XML format. By contrast, `GetHTML` expects the input document to be in HTML format and returns a modified HTML document in which the recognized terms have been marked up by HTML tags.

By default the tagger will auto detect the organisms mentioned in a document and subsequently tag their proteins/gene. If a user is interested in genes/proteins of specific organisms, tagging of them can be forced via the `entity_types` parameter (e.g. `entity_types=9606` will force tagging of human proteins). Unless `auto_detect` is disabled (`auto_detect=0`), the tagger will also tag genes/proteins from any organisms that are explicitly mentioned in the text. If a user is not interested in the identification of genes/proteins, auto detection should be disabled.

For further documentation on the API of the real-time tagger, please refer to O’Donoghue *et al.* (2) and http://extract.hcmr.gr/ (purple section of the help page).

The pre-tagged PubMed abstracts can also be queried via HTTP requests, using this syntax: http://{resource}.jensenlab.org/document/{pmid}/annotations/, where {`resource`} is either `compartments, tissues, diseases,` or `organisms` and {`pmid`} is the PubMed ID of the abstract of interest. For more general information on the RESTful Open Annotation API, please refer to Pyysalo *et al.* (19) and https://github.com/restful-open-annotation/.

## Future plans

The core tagger obviously has the full information about which entities were found where in the text. However, this level of detail is not currently exposed by the REST API. Our plan for the hackathon is to address this shortcoming in a manner that makes the real-time tagger provide automatic annotations according to the PubAnnotation (20, 21) standard and, if time permits, also the the Open Annotation (19) standard. In the case of the pre-tagged corpora, we plan to make these available also in PubAnnotation format.

## Acknowledgments

This work was funded by the LifeWatchGreece Research Infrastructure (384676-94/GSRT/NSRF(C&E)) and the Novo Nordisk Foundation (NNF14CC0001).

## References

1. Pafilis E, O’Donoghue SI, Jensen LJ, et al. (2009). Reflect: augmented browsing for the life scientist. Nat. Biotechnol., 27:508–510.

2. O’Donoghue SI, Horn H, Pafilis E, et al. (2010). Reflect: A practical approach to web semantics. Journal of Web Semantics, 8:182–189.

3. Pafilis E, Buttigieg PL, Ferrell B, et al. (2015). EXTRACT: Interactive extraction of environment metadata and term suggestion for metagenomic sample annotation. Proceedings of the Fifth BioCreative Challenge Evaluation Workshop, 384–395.

4. Pafilis E, Pletscher-Frankild S, Fanini L, et al. (2013). The SPECIES and ORGANISMS resources for fast and accurate identification of taxonomic names in text. PLoS One, 8:e65390.

5. Pafilis E, Pletscher-Frankild S, Schnetzer J, et al. (2015). ENVIRONMENTS and EOL: identification of Environment Ontology terms in text and the annotation of the Encyclopedia of Life. Bioinformatics, 31:1872–1874.

6. Szklarczyk D, Franceschini A, Wyder S, et al. (2015). STRING v10: protein-protein interaction networks, integrated over the tree of life. Nucleic Acids Res., 43:D447–D452.

7. Kuhn M, Szklarczyk D, Pletscher-Frankild S, et al. (2014). STITCH 4: integration of protein-chemical interactions with user data. Nucleic Acids Res., 42:D401–D407.

8. Binder JX, Pletscher-Frankild S, Tsafou K, et al. (2014). COMPARTMENTS: unification and visualization of protein subcellular localization evidence. Database, 2014:bau012.

9. Santos A, Tsafou K, Stolte C, et al. (2015). Comprehensive comparison of large-scale tissue expression datasets. PeerJ, 3:e1054.

10. Pletscher-Frankild S, Pallejà A, Tsafou K, et al. (2015). DISEASES: text mining and data integration of disease-gene associations. Methods, 74:83–89.

11. Cunningham F, Amode MR, Barrell D, et al. (2015). Ensembl 2015. Nucleic Acids Res., 43:D662–D669.

12. Tatusova T, Ciufo S, Fedorov B, et al. (2014). RefSeq microbial genomes database: new representation and annotation strategy. Nucleic Acids Res., 42:D553–D559

13. Bolton E, Wang Y, Thiessen PA and Bryant SH (2008). PubChem: integrated platform of small molecules and biological activities. Annu. Rep. Comput. Chem., 4:217–241.

14. Federhen S (2015). Type material in the NCBI Taxonomy Database. Nucleic Acids Res., 43:D1086–D1098.

15. Ashburner M, Ball CA, Blake JA, et al. (2000). Gene ontology: tool for the unification of biology. The Gene Ontology Consortium. Nat. Genet., 25:25–29.

16. Chang A, Schomburg I, Placzek S, et al. (2015). BRENDA in 2015: exciting developments in its 25th year of existence. Nucleic Acids Res., 43:D439–D446.

17. Kibbe WA, Arze C, Felix V, et al. (2015). Disease Ontology 2015 update: an expanded and updated database of human diseases for linking biomedical knowledge through disease data. Nucleic Acids Res., 43:D1071–D1078.

18. Buttigieg PL, Morrison N, Smith B, et al. (2013) The environment ontology: contextualising biological and biomedical entities. J. Biomed. Semant., 4:43.

19. Pyysalo S, Campos J, Cejuela JM, et al. (2015). Sharing annotations better: RESTful Open Annotation. Proceedings of ACL-IJCNLP 2015, 91–96.

20. Kim JD and Wang Y (2012) PubAnnotation: a persistent and sharable corpus and annotation repository. Proceedings of BioNLP 2012, 202–205.

21. ￼Kim JD, Cohen KB and Kim JJ (2015). PubAnnotation-query: a search tool for corpora with multilayers of annotation. BMC Proceedings, 9(Suppl 5):A3.

